# Prediction, scanning and designing of TNF-α inducing epitopes for human and mouse

**DOI:** 10.1101/2022.08.02.502430

**Authors:** Anjali Dhall, Sumeet Patiyal, Shubham Choudhury, Shipra Jain, Kashish Narang, Gajendra P. S. Raghava

## Abstract

Tumor Necrosis Factor alpha (TNF-α) is a pleiotropic pro-inflammatory cytokine that plays a crucial role in controlling signaling pathways within the immune cells. Recent studies reported that the higher expression levels of TNF-α is associated with the progression of several diseases including cancers, cytokine release syndrome in COVID-19 and autoimmune disorders. Thus, it is the need of the hour to develop immunotherapies or subunit vaccines to manage TNF-α progression in various disease conditions. In the pilot study, we have proposed a host-specific in-silico tool for the prediction, designing and scanning of TNF-α inducing epitopes. The prediction models were trained and validated on the experimentally validated TNF-α inducing/non-inducing for human and mouse hosts. Firstly, we developed alignment free (machine learning based models using composition of peptides) methods for predicting TNF-α inducing peptides and achieved maximum AUROC of 0.79 and 0.74 for human and mouse hosts, respectively. Secondly, alignment based (using BLAST) method has been used for predicting TNF-α inducing epitopes. Finally, a hybrid method (combination of alignment free and alignment-based method) has been developed for predicting epitopes. Our hybrid method achieved maximum AUROC of 0.83 and 0.77 on an independent dataset for human and mouse hosts, respectively. We have also identified the potential TNF-α inducing peptides in different proteins of HIV-1, HIV-2, SARS-CoV-2 and human insulin. Best models developed in this study has been incorporated in a webserver TNFepitope (https://webs.iiitd.edu.in/raghava/tnfepitope/), standalone package and GitLab (https://gitlab.com/raghavalab/tnfepitope).

**Key Points:**

- TNF-α is a multifunctional pleiotropic pro-inflammatory cytokine.
- Anti-TNF-α therapy used as an effective treatment in several autoimmune disorders.
- Composition-based features generated using Pfeature for each peptide sequence.
- Alignment-based and alignment-free models developed.
- Prediction and scanning of TNF-α inducing regions in antigens.
- TNFepitope is available as a web-server, standalone package and GitLab.

## Introduction

Tumor Necrosis Factor alpha (TNF-α), is a classical, pleiotropic pro-inflammatory cytokine that function by promoting cellular signal activation and trafficking of leukocytes to inflammatory sites [1]. During acute inflammation, TNF-α cytokine is released by macrophages/monocytes or via other cell types (e.g., B cells, T cells, mast cells, fibroblasts), which further regulates haematopoiesis, immune responses, tumor regression and various infections [2–6]. TNF-α is the first “adipokine” reported in literature to be produced from adipose tissue [7–9]. It plays a significant role in various biological processes, including immunomodulation, fever, inflammatory response, inhibition of tumor formation, and inhibition of virus replication [10]. In its active form TNF-α molecule exists as a homotrimer, where it binds to homotrimeric TNFRs receptors to induce signaling [11]. Most of the downstream functions of TNF-α are executed via binding with two distinct receptors: TNFR1 and TNFR2 [11]. Pleiotropic biological effects of TNF-α are based on the interactions between TNF and its receptors (both circulating and membrane-bound) [3]. Binding of TNF-α to its receptor can initiate several signaling pathways, including the activation of transcription factors (e.g., nuclear factor-κB [NF-κB]), protein kinases (e.g., c-Jun N-terminal kinase [JNK], p38 MAP kinase), and proteases (e.g., caspases) that markedly impact immune and inflammatory responses [12].

Recent studies revealed that TNF-α is involved in various physiological effects such as induction of pro-inflammatory interleukins (IL-1 and IL-6) [13–15]. Studies also shows that TNF-α and IL-1β have been found to be implicated in the pathogenesis of myocardial dysfunction in ischemia-reperfusion injury, sepsis, chronic heart failure, viral myocarditis, and cardiac allograft rejection [16–18]. In addition, TNF-α also interacts with various cytokines/chemokines and regulates signaling pathways in various other disease states [19]. For example, Guo et. al., reported that cytokine release syndrome in COVID-19 patients is associated with the increased levels of TNF-α, IL-6, IL-2, IL-7, and IL-10 cytokines [20]. In addition, a number of studies reported the direct relationship of TNF-α and IL-6 cytokines in the severity and survival of COVID-19 patients [21–23]. Therefore, several anti-TNF inhibitors are available in the market which can block the over production of TNF-α in different disease conditions. In literature, studies have reported wide use of anti-TNF therapy for effective treatment of rheumatoid arthritis (RA), spondyloarthropathy, psoriasis and inflammatory bowel disease [24–27]. In the recent times, anti-TNF-α therapy has reported beneficial effects by not only restoring aberrant TNF-mediated immune mechanisms, but also by de-activating pathogenic fibroblast-like mesenchymal cells [28].

As reported in literature, TNF-α is a key cytokine involved in several diseases and their increasing severity. Therefore, it can act as a primary target cytokine in disease progression. This creates a need to develop a computational tool, for predicting TNF-α inducing peptides using sequence information. In present study, we have come up with an in-silico method to classify the TNF-α inducing and non-inducing epitopes. We have developed this tool using experimentally validated TNF-α inducing and non-inducing peptides for human and mouse hosts. In addition, we have used randomly generated peptides from SwissProt database [29]. We have developed prediction models using various machine learning classifiers and evaluated performance on the independent dataset.

## Material and methods

### Overall Workflow

The complete workflow of the current study is illustrated in Figure 1.

**Figure 1:**
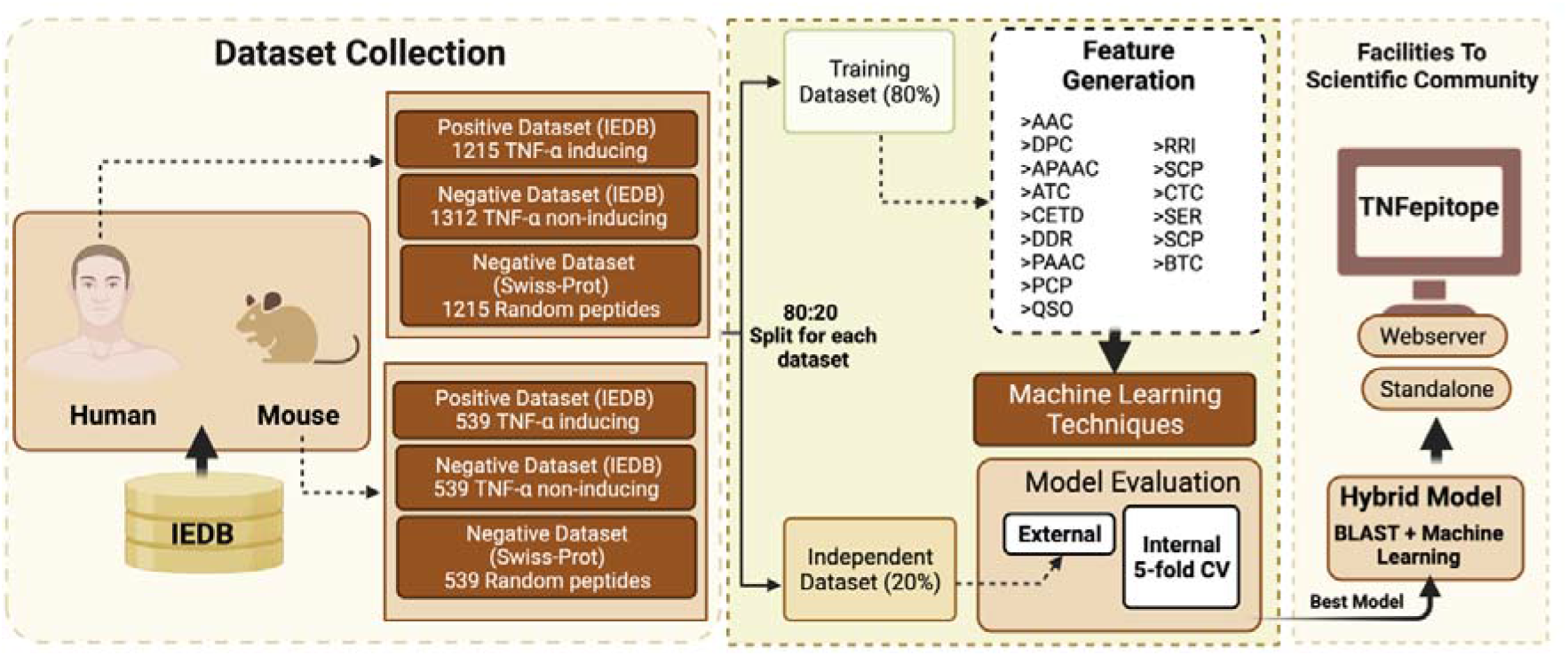
Overall architecture of the study; including data creation, feature generation, model building and webserver.

### Dataset collection and preprocessing

We have collected 3635 TNF-α inducing peptides/epitopes from the immune epitope database (IEDB [30]). At first, we filtered the dataset based on the hosts and observed that 3177 peptides are experimentally validated on human or mouse hosts, and only a few epitopes are available for other hosts. So, we selected only two major hosts (i.e., human and mouse). We have checked the length distribution of epitopes and observed that most of the peptides belong to the range of 8-20 amino-acid residues. After removing the redundancy, we obtained 1215 and 539 TNF-α inducing epitopes for humans and mouse, respectively.

In this study, we have two separate negative datasets for both human and mouse. The first negative dataset was collected from IEDB, containing 2383 experimentally validated TNF-α non-inducing epitopes for both the hosts. After preprocessing, we obtain 1312 unique TNF-α non-inducing epitopes with a range of (8-20 amino-acids) in the case of human host. On the other side, we have 539 unique TNF-α non-inducing epitopes for the mouse with 8-20 aminoacids residues range. Finally, the main dataset for human incorporates 1215 TNF-α inducing and 1312 TNF-α non-inducing peptides. On the other side, in case of mouse we obtain a total of 539 TNF-α inducing and 539 non-inducing peptides in the main dataset. The second negative dataset was created using the Swiss-Prot database [29]. Here, we have generated random peptides for human and mouse to construct another negative dataset. Finally, the alternate dataset for human incorporates 1215 TNF-α inducing and 1215 randomly generated peptides sing Swiss-Prot database. Similarly, in case of mouse we get a total of 539 TNF-α inducing and 539 randomly generated peptides. After generating the final datasets for human and mouse hosts; each dataset was divided into training and independent/validation set. Here, the complete dataset splitted into 80:20 ratio where 80% data was used to train the models and 20% data was used for the validation purpose.

### Composition-based analysis

We have used Pfeature [31] to calculate the amino acid composition (AAC) of main and alternate datasets. Using the compositional analysis, we understand the similarity between the different peptide sequences taken from positive and negative datasets. Using the following equation 1, we have generated a feature vector of length 20, which specify the percent composition of 20 amino-acid residues.

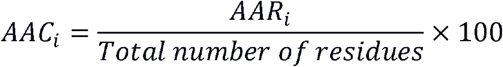

where AAC_i_ and AAR_i_ are the percentage composition and number of residues of type i in a peptide, respectively.

### WebLogo

We have used WebLogo (http://weblogo.threeplusone.com) [32] in order to generate sequence logos of TNF-α inducing epitopes. Here x-axis represents the amino-acid residues and y-axis presents the bit-score which shows the importance of particular residue at a given position. WebLogo takes a fixed length vector of input peptide sequences. In order to create a fix length vector, we have considered eight amino-acids from the N-terminal and eight-residues from the C-terminal, as eight is the minimum length of eptides in our dataset and merged them to generated a fixed length vector of sixteen residues for both human and mouse TNF-α inducing epitopes.

### Feature generation

In the current study, we have calculated a wide range of features using the sequence information of peptide sequences. We have used Pfeature [31] standalone package in order to calculate the composition-based features for our datasets. We have computed a total of 1163 features for each epitope/peptide sequence in both positive and negative datasets. We have computed twelve different types of descriptors/features such as AAC (Amino acid composition), DPC (Di-peptide composition), APAAC (Amphiphilic pseudo amino acid composition), ATC (Atomic composition), CETD (Composition-enhanced transition distribution), DDR (Distance distribution of residue), PAAC (Pseudo amino acid composition), PCP (Physico-chemical properties composition), QSO (Quasi-sequence order), RRI (Residue repeat Information), SPC (Shannon entropy of physico-chemical properties), CTC (Conjoint triad descriptors), etc. In this study, we have developed prediction models using each feature as well as combining all the features.

### Machine learning and Cross-validation Techniques

In order to develop the prediction models, we have used various machine learning algorithms such as Random Forest (RF), Decision Tree (DT), Gaussian Naive Bayes (GNB), Logistic Regression (LR), Support Vector Classifier (SVC), K-Nearest Neighbor (KNN) and Extra Tree (ET). We have trained the parameters on training dataset and predictions were made on the independent dataset. Scikit-learn [33] python library was used in the study for the implementation of various classifiers. We have employed five-fold cross validation technique in order to evade the curse of biasness and overfitting. In the five-fold cross-validation, first the training dataset was divided into five equal sets; where four sets were used for training and fifth set was used for testing. This process is repeated five times where each part gets utilized for testing of the model as shown in some previous studies [34–40]. Of note, the final performance is the mean of the performance resulted after each iteration.

### Similarity Search Method

We have used BLAST [41] to implement similarity search or alignment-based approach; where we classify the epitopes as TNF-α inducing and non-inducing on the basis of the similarity. Here, we have used NCBI-BLAST+ version 2.2.29 (blastp suite) for similarity search and makeblastdb suite of NCBI-BLAST+ for the creation of custom database. We have created a custom database using the training dataset; and sequences of validation dataset were searched against the created database. Based on the hits and their similarity with the customized database, we assign class as TNF-α inducer or non-inducer. Currently we have considered only top-hit of BLAST (i.e., if the top-hit of BLAST is against the TNF-α inducer peptide then the query sequence was assigned as TNF-α inducing peptide or vice-versa). To identify the optimal value of e-value; we run the BLAST at various e-values cut-offs varying from 1e-6 to 1e+3.

### Hybrid Model

In order to improve the prediction, we have applied the hybrid approach in which we merge alignment-based (BLAST) and alignment-free (machine learning based prediction). Here, first we classify the peptide/epitope based on the BLAST. After that, we add ‘+0.5’ score for the correct positive prediction i.e., TNF-α inducing peptide, ‘-0.5’ score integrated for the negative predictions i.e., TNF-α non-inducing peptide and ‘0’ score if no-hit was found. Further, we incorporate the prediction score calculated using machine learning based models. Finally, we combine the BLAST score and machine learning prediction score to make final predictions.

### Performance Evaluation

The performance of different models were evaluated using standard performance evaluation parameters sensitivity, specificity, accuracy, Area Under Receiver Operating Characteristics (AUROC) curve, Area Under the Precision-Recall Curve (AUPRC), Matthews Correlation Coefficient (MCC), and F1-score. We have computed both threshold-dependent (including sensitivity, specificity, accuracy, F1-score, and MCC) and independent parameters such as AUROC and AUPRC. The equations of evaluation parameters is provided in equations (2-6).

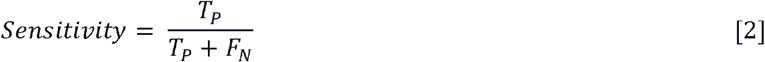

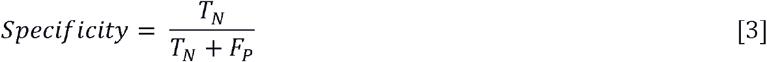

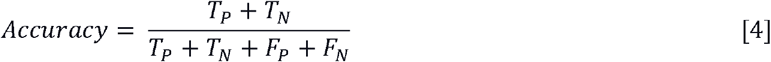

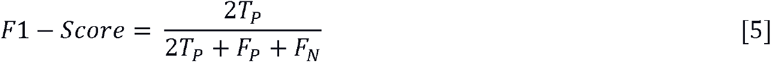

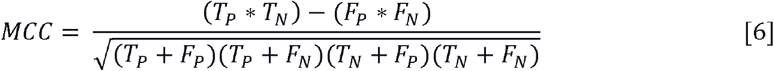

Where, FP is false positive, FN is false negative, TP is true positive and TN is true negative.

## Results

### Compositional Analysis

We have computed amino acid composition for the main and alternate datasets for human and mouse hosts. After that, we have calculated the average compositions of TNF-α inducing and non-inducing peptides. As depicted in Figure 2A, in case of human dataset amino acids such as leucine (L), valine (V), tyrosine (Y), and tryptophan (W) having higher composition in the TNF-α inducing peptides in comparison with the TNF-α non-inducing and random peptides. Similarly, the average composition of residues like alanine (A), isoleucine (I), asparagine (N), and serine (S) are more abundant in TNF-α inducing peptides of mouse dataset (See Figure 2B).

**Figure 2:**
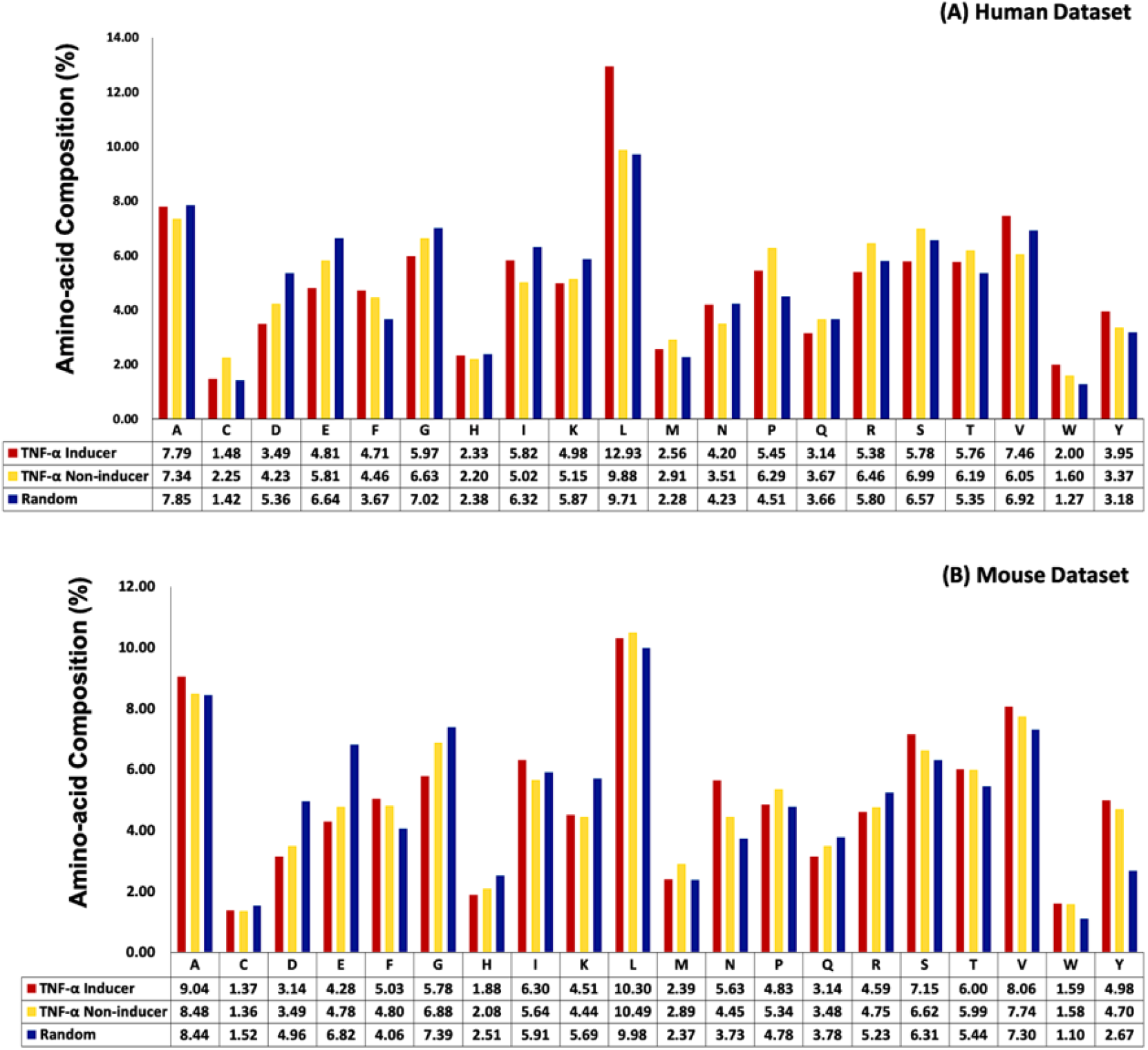
Average amino-acid composition of TNF-α inducing, TNF-α non-inducing and random peptides.

### Positional Conservation Analysis

In this analysis, we study the preference of residues at particular position in the TNF-α inducing epitopes for human and mouse dataset. In the case of human TNF-α inducing epitopes, residues ‘L’ is highly conserved at most of the positions, whereas ‘V’ is preferred at 9^th^ and 16^th^ positions; ‘A’ is located on 7^th^, 9^th^, 10^th^, 11^th^, 12^th^, 13^th^ and 16^th^ positions (See Figure 3A). In the case of mouse, TNF-α inducing epitopes ‘L’ is highly dominated on 2^nd^, 3^rd^, 8^th^, 9^th^, 12^th^, 13^th^ and 16^th^ positions; similarly residue ‘N’ is highly conserved at 5^th^ and 13^th^ positions; however, ‘A’ is predominated on 5^th^, 8^th^, 9^th^, 13^th^, 16^th^ positions, as shown in Figure 3B.

**Figure 3:**
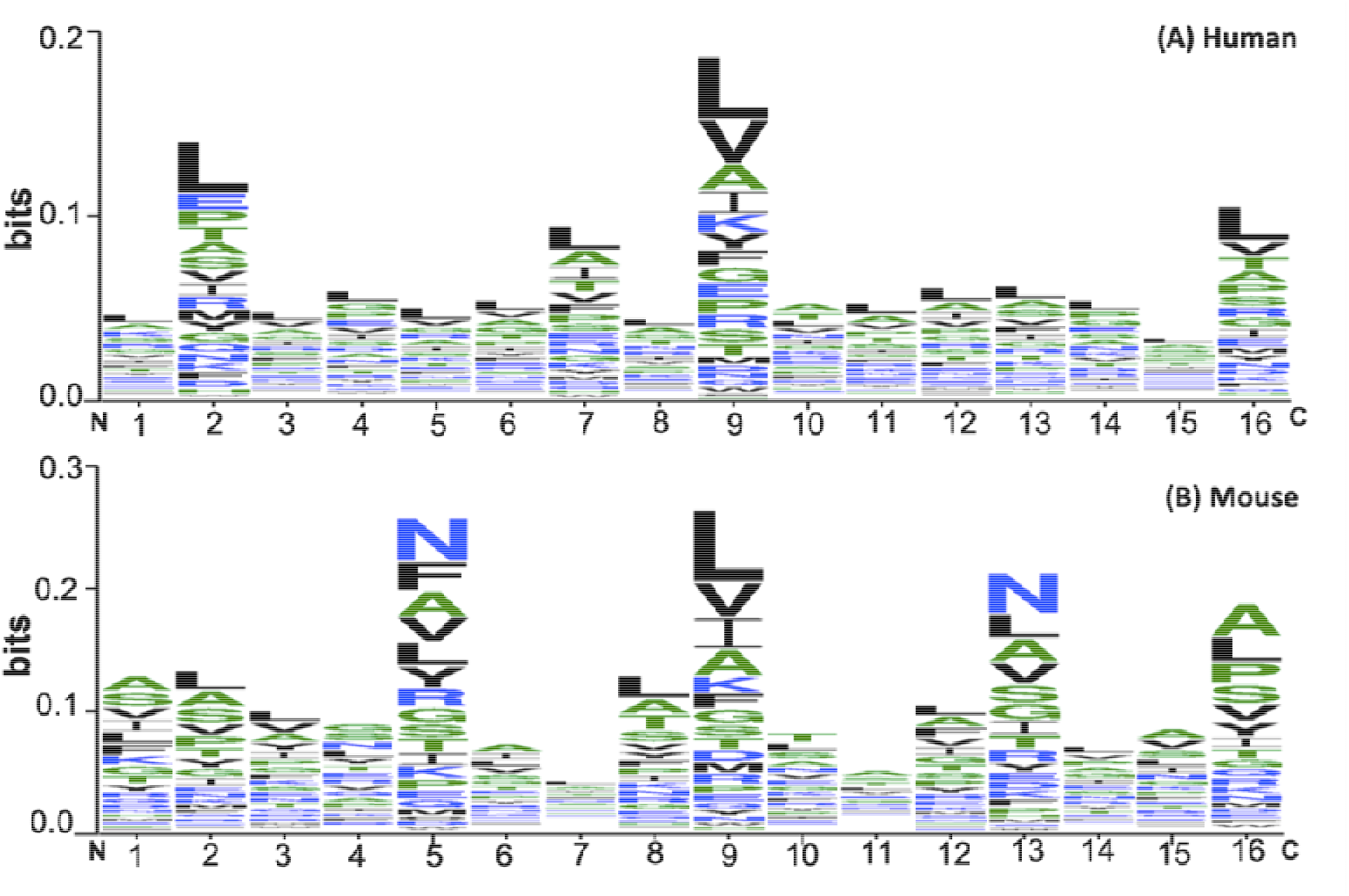
WebLogo representation of TNF-α inducing peptides in human and mouse datasets.

### Machine Learning based Predictions

We have developed prediction models using different classifiers such as DT, RF, GNB, KNN, SVC, LR and ET on main and alternate datasets of both human and mouse hosts. For this we have generated 15 different types of composition based features using Pfeature standalone. We evaluated the performance on different features as well as combining all the features.

### Performance of Composition-based Features

Here, we have computed performance on 15 different features. We have observed that RF and ET classifiers performed best among the other classifiers (See Supplementary Table S1). As shown in Table 2, in the case of human host, we achieved maximum performance on main dataset with an AUROC of 0.79, MCC of 0.45 on the independent dataset using DPC based features. APAAC and SER based features also performed quite well on independent dataset with an AUROC of 0.78 and AUPRC of 0.75. In the case of alternate dataset we attains maximum AUROC of 0.71, AUPRC of 0.73 and MCC of 0.31 using DPC based features. While combining all the features we are getting (0.77 and 0.71) AUROC on main and alternate dataset, respectively. Other composition-based features, perform poor on both main and alternate dataset. The complete results of all the classifiers for the features are shown in Supplementary Table S2.

**Table 1:**
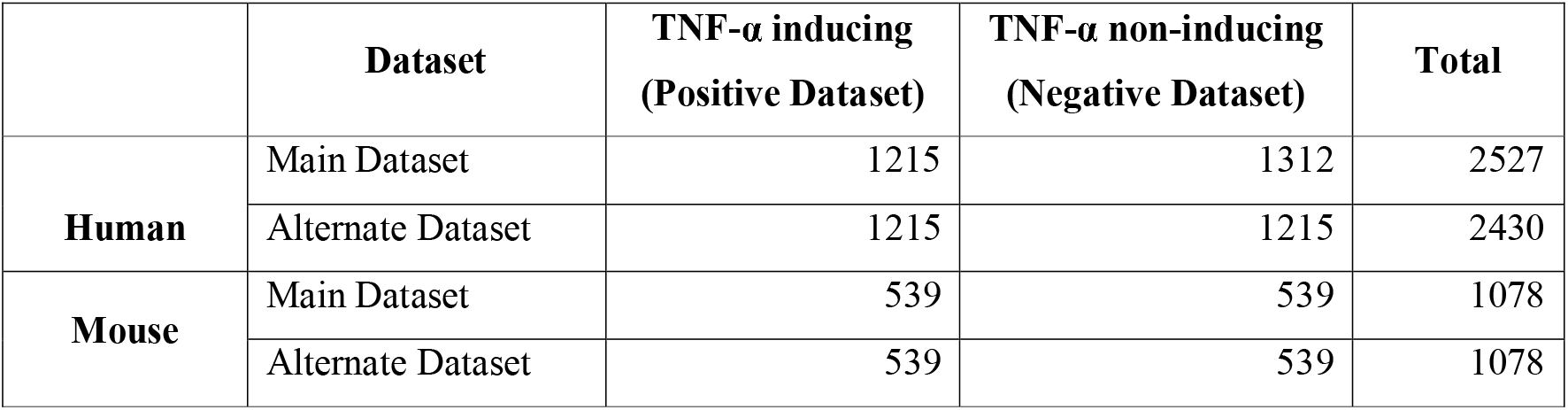
Distribution of TNF-α inducing and non-inducing peptides extracted from IEDB and Swiss-Prot database.

**Table 2:** Performance of independent dataset developed using 15 types of composition-based features for human main and alternate datasets.

| Feature Type | Main Dataset |  |  |  |  |  | Alternate Dataset |  |  |  |  |  |
| --- | --- | --- | --- | --- | --- | --- | --- | --- | --- | --- | --- | --- |
|  | Sens | Spec | Acc | AUROC | AUPRC | MCC | Sens | Spec | Acc | AUROC | AUPRC | MCC |
| AAC | 55.97 | 58.56 | 57.31 | 0.63 | 0.61 | 0.15 | 63.37 | 66.26 | 64.82 | 0.70 | 0.72 | 0.30 |
| DPC | 72.02 | 72.62 | 72.33 | 0.79 | 0.76 | 0.45 | 68.72 | 61.73 | 65.23 | 0.71 | 0.73 | 0.31 |
| ATC | 55.97 | 58.56 | 57.31 | 0.63 | 0.61 | 0.15 | 59.67 | 58.03 | 58.85 | 0.61 | 0.62 | 0.18 |
| APAAC | 68.31 | 74.91 | 71.74 | 0.78 | 0.75 | 0.43 | 63.37 | 67.49 | 65.43 | 0.70 | 0.73 | 0.31 |
| BTC | 69.55 | 68.82 | 69.17 | 0.69 | 0.64 | 0.38 | 55.97 | 50.62 | 53.29 | 0.55 | 0.53 | 0.07 |
| CETD | 66.67 | 70.34 | 68.58 | 0.74 | 0.72 | 0.37 | 61.32 | 61.32 | 61.32 | 0.64 | 0.64 | 0.23 |
| CTD | 61.32 | 66.92 | 64.23 | 0.70 | 0.65 | 0.28 | 62.14 | 61.73 | 61.93 | 0.66 | 0.68 | 0.24 |
| DDR | 72.02 | 73.76 | 72.93 | 0.77 | 0.74 | 0.46 | 62.55 | 64.61 | 63.58 | 0.70 | 0.71 | 0.27 |
| PAAC | 68.31 | 74.14 | 71.34 | 0.78 | 0.75 | 0.43 | 65.02 | 65.43 | 65.23 | 0.70 | 0.72 | 0.31 |
| PCP | 64.61 | 67.68 | 66.21 | 0.73 | 0.72 | 0.32 | 62.96 | 63.37 | 63.17 | 0.67 | 0.67 | 0.26 |
| QSO | 62.55 | 71.86 | 67.39 | 0.72 | 0.71 | 0.35 | 63.79 | 65.43 | 64.61 | 0.69 | 0.71 | 0.29 |
| RRI | 62.55 | 68.06 | 65.42 | 0.73 | 0.70 | 0.31 | 62.96 | 57.20 | 60.08 | 0.66 | 0.69 | 0.20 |
| SEP | 63.37 | 60.84 | 62.06 | 0.69 | 0.67 | 0.24 | 43.62 | 57.61 | 50.62 | 0.51 | 0.50 | 0.01 |
| SER | 67.08 | 73.38 | 70.36 | 0.78 | 0.75 | 0.41 | 64.61 | 67.90 | 66.26 | 0.70 | 0.73 | 0.33 |
| SCP | 66.67 | 73.38 | 70.16 | 0.74 | 0.73 | 0.40 | 65.02 | 62.14 | 63.58 | 0.68 | 0.70 | 0.27 |
| ALL_COMP | 68.31 | 74.91 | 71.73 | 0.77 | 0.74 | 0.433 | 65.43 | 65.02 | 65.22 | 0.71 | 0.73 | 0.30 |

In case of mouse dataset, RF-based classifier perform well with an AUROC of 0.74, AUPRC of 0.76 and MCC of 0.34 on alternate dataset using DPC as input feature (See Table 3). Similarly, we achieved an equivalent performance (i.e., AUROC = 0.72, MCC = 0.30, and AUPRC = 0.73) using AAC based features on the alternate dataset. In addition, RRI, DDR and APAAC also perform quite well with AUROC>0.72 on alternate dataset. However, the performance of machine learning models is comparatively low on the main dataset. The complete results on training and independent dataset is provide in Supplementary Table S3, S4.

### Performance of Hybrid Models

In this study, we have developed a hybrid model to classify TNF-α inducing and non-inducing peptides. At first, we have used the similarity search approach (BLAST) for the prediction of positive and negative peptides. As shown in Table 2 and 3, DPC based features outperformed on both human and mouse prediction models. Hence, we combined BLAST similarity scores and machine learning scores computed using DPC features to make the final predictions. As shown in Supplementary Table S2, RF and ET based models performed well on main and alternate human datasets, respectively. We have used DPC features and best models to calculate the performance of hybrid models at different e-value cutoffs on independent datasets as exhibit in Table 4 for human host. We obtained highest performance at e-value (1.00E-01) with AUROC of (0.83 and 0.79), AUPRC of (0.80 and 0.84), MCC of (0.52 and 0.41) on main and alternate dataset, respectively (See Table 4). The complete results of training and independent datasets are provided in Supplementary Table S3.

**Table 3:** Performance of independent dataset developed using 15 types of composition-based features for mouse main and alternate datasets.

| Feature Type | Main Dataset |  |  |  |  |  | Alternate Dataset |  |  |  |  |  |
| --- | --- | --- | --- | --- | --- | --- | --- | --- | --- | --- | --- | --- |
|  | Sens | Spec | Acc | AUROC | AUPRC | MCC | Sens | Spec | Acc | AUROC | AUPRC | MCC |
| AAC | 62.18 | 60.56 | 61.37 | 0.67 | 0.66 | 0.23 | 64.82 | 64.82 | 64.82 | 0.72 | 0.73 | 0.30 |
| DPC | 58.47 | 59.86 | 59.17 | 0.63 | 0.62 | 0.18 | 66.67 | 67.59 | 67.13 | 0.74 | 0.76 | 0.34 |
| ATC | 51.97 | 50.35 | 51.16 | 0.54 | 0.53 | 0.02 | 55.56 | 62.04 | 58.80 | 0.65 | 0.62 | 0.18 |
| APAAC | 62.18 | 60.09 | 61.14 | 0.65 | 0.63 | 0.22 | 63.89 | 65.74 | 64.82 | 0.72 | 0.73 | 0.30 |
| BTC | 51.51 | 52.44 | 51.97 | 0.55 | 0.53 | 0.04 | 51.85 | 58.33 | 55.09 | 0.56 | 0.55 | 0.10 |
| CETD | 56.15 | 58.24 | 57.19 | 0.62 | 0.63 | 0.14 | 63.89 | 66.67 | 65.28 | 0.70 | 0.73 | 0.31 |
| CTD | 51.51 | 53.13 | 52.32 | 0.56 | 0.57 | 0.05 | 65.74 | 63.89 | 64.82 | 0.68 | 0.68 | 0.30 |
| DDR | 56.85 | 59.86 | 58.35 | 0.62 | 0.63 | 0.17 | 69.44 | 67.59 | 68.52 | 0.74 | 0.75 | 0.37 |
| PAAC | 60.79 | 61.02 | 60.91 | 0.65 | 0.64 | 0.22 | 67.59 | 65.74 | 66.67 | 0.72 | 0.73 | 0.33 |
| PCP | 57.77 | 61.49 | 59.63 | 0.61 | 0.59 | 0.19 | 56.48 | 69.44 | 62.96 | 0.70 | 0.70 | 0.26 |
| QSO | 58.01 | 58.47 | 58.24 | 0.60 | 0.59 | 0.17 | 61.11 | 70.37 | 65.74 | 0.73 | 0.74 | 0.32 |
| RRI | 59.86 | 60.79 | 60.33 | 0.63 | 0.62 | 0.21 | 65.74 | 66.67 | 66.20 | 0.75 | 0.74 | 0.32 |
| SEP | 55.68 | 54.06 | 54.87 | 0.57 | 0.56 | 0.10 | 36.11 | 51.85 | 43.98 | 0.45 | 0.46 | -0.12 |
| SER | 60.56 | 62.41 | 61.49 | 0.67 | 0.66 | 0.23 | 67.59 | 69.44 | 68.52 | 0.73 | 0.74 | 0.37 |
| SCP | 57.77 | 58.47 | 58.12 | 0.61 | 0.59 | 0.16 | 60.19 | 69.44 | 64.82 | 0.69 | 0.66 | 0.30 |
| ALL_COMP | 62.96 | 62.96 | 62.96 | 0.67 | 0.67 | 0.26 | 64.81 | 68.51 | 66.67 | 0.73 | 0.73 | 0.33 |

**Table 4:** Performance of hybrid model on human main and alternative independent datasets.

| E-value | Main Dataset |  |  |  |  |  | Alternate Dataset |  |  |  |  |  |
| --- | --- | --- | --- | --- | --- | --- | --- | --- | --- | --- | --- | --- |
|  | Sens | Spec | Acc | AUROC | AUPRC | MCC | Sens | Spec | Acc | AUROC | AUPRC | MCC |
| 1.00E-06 | 72.43 | 76.34 | 74.46 | 0.82 | 0.79 | 0.49 | 65.02 | 65.02 | 65.02 | 0.72 | 0.76 | 0.30 |
| 1.00E-05 | 73.66 | 77.48 | 75.64 | 0.81 | 0.77 | 0.51 | 67.49 | 65.84 | 66.67 | 0.73 | 0.77 | 0.33 |
| 1.00E-04 | 72.84 | 75.57 | 74.26 | 0.81 | 0.76 | 0.48 | 66.26 | 69.14 | 67.70 | 0.73 | 0.77 | 0.35 |
| 1.00E-03 | 72.43 | 77.10 | 74.85 | 0.81 | 0.77 | 0.50 | 65.02 | 69.14 | 67.08 | 0.73 | 0.78 | 0.34 |
| 1.00E-02 | 74.90 | 76.72 | 75.84 | 0.82 | 0.77 | 0.52 | 68.72 | 69.55 | 69.14 | 0.78 | 0.83 | 0.38 |
| 1.00E-01 | 76.13 | 75.95 | 76.04 | 0.83 | 0.80 | 0.52 | 70.37 | 70.78 | 70.58 | 0.79 | 0.84 | 0.41 |
| 1.00E+00 | 76.54 | 75.95 | 76.24 | 0.83 | 0.81 | 0.53 | 68.72 | 67.90 | 68.31 | 0.77 | 0.81 | 0.37 |
| 1.00E+01 | 73.25 | 74.81 | 74.06 | 0.82 | 0.79 | 0.48 | 67.49 | 68.31 | 67.90 | 0.74 | 0.78 | 0.36 |
| 1.00E+02 | 72.84 | 72.14 | 72.48 | 0.82 | 0.79 | 0.45 | 67.08 | 67.49 | 67.28 | 0.73 | 0.78 | 0.35 |

Besides this, we have applied similar approach on mouse dataset, as provided in Supplementary Table S4, RF based model outperforms the other classifier on both main and alternate human datasets with DPC based features. Using hybrid model, we achieved highest performance at e-value (1.00E-01) with AUROC of (0.70 and 0.77), AUPRC of (0.69 and 0.81), MCC of (0.28 and 0.34) on main and alternate dataset, respectively. The comprehensive results of training and independent datasets are given in Supplementary Table S5.

### Services to Scientific Community

We have developed a web-server named ‘TNFepitope’ for the prediction of TNF-α inducing and non-inducing epitopes using sequence information. The best prediction models for human and mouse hosts were integrated in the webserver. We have incorporated five major modules in the server (i) Predict; (ii) Design; (iii) Scan; (iv) Blast Search; and (v) Standalone. ‘Predict’ module facilitates the users to classify TNF-α inducing peptides from non-inducing peptides. The ‘Design’ module provide facility to the user to design/create all possible mutants of query sequence and predict if that can induce the TNF-α release. The ‘Scan’ module allows the user to map/scan the TNF-α secretion portion in the given protein sequence. The ‘BLAST Search’ module entirely based on similarity search algorithm, the input sequence is hit against the customized database created using the known TNF-α inducing and non-inducing peptides. The submitted amino-acid sequence is predicted as TNF-α inducer/non-inducer based on the similarity. ‘TNFepitope’ server was developed using HTML, JAVA and PHP scripts; it is compatible with a number of devices such as laptops, iPhone, phones, etc. The webserver (https://webs.iiitd.edu.in/raghava/tnfepitope), standalone package (https://webs.iiitd.edu.in/raghava/tnfepitope/package.php) and GitLab (https://gitlab.com/raghavalab/tnfepitope) are freely-accessible. Figure 4 depicts all the major modules of TNFepitope webserver.

**Figure 4:**
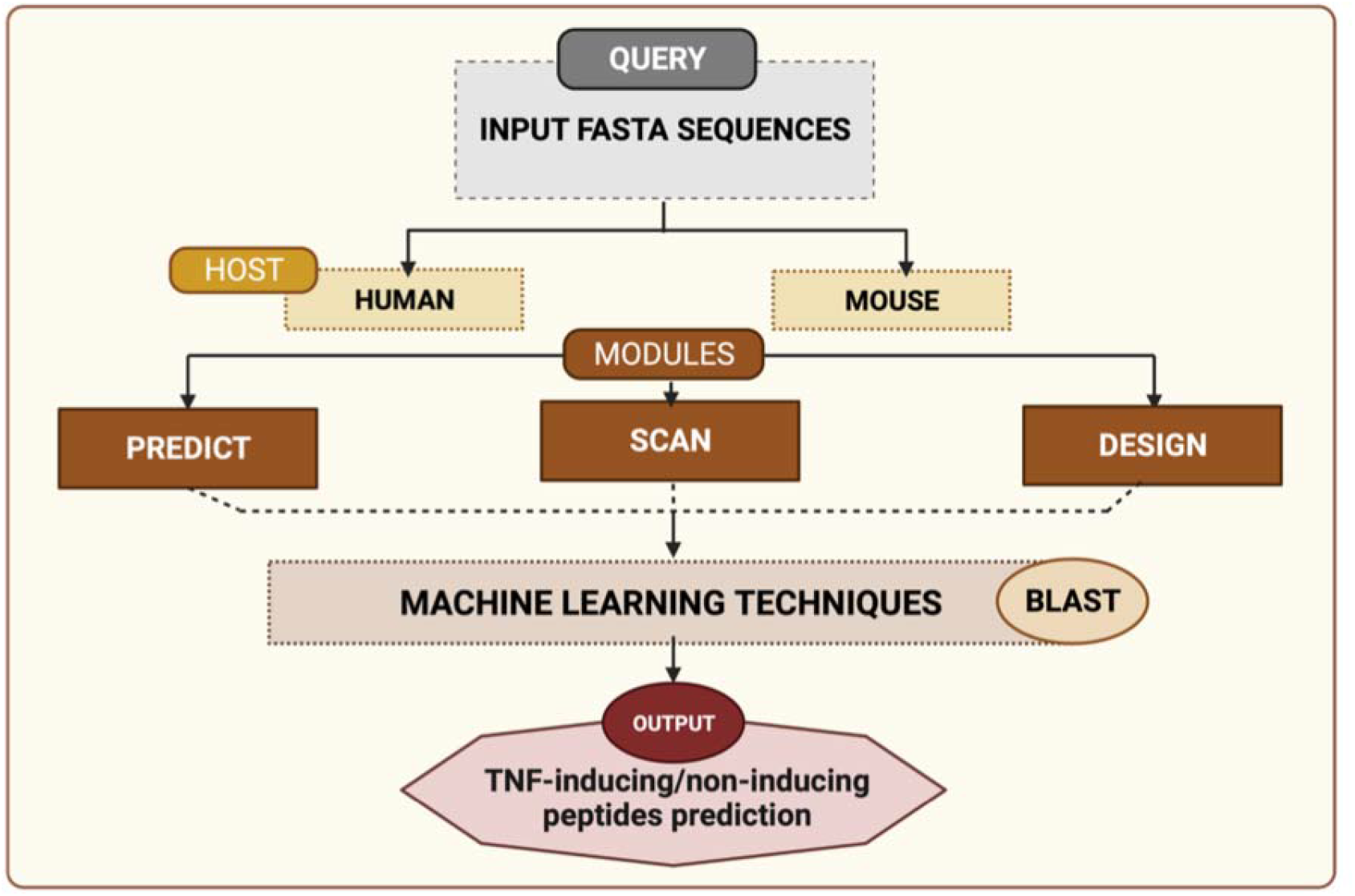
Schematic representation of different modules of TNFepitope server.

### Case Study

In order to demonstrate the application of our work, we predicted TNF-α inducing epitopes using ‘Scan’ module of TNFepitope webserver with default parameters (i.e., length of peptide 15 and threshold 0.45 with the hybrid method). Here, we have used three viral proteins (envelope glycoprotein of HIV-1, HIV-2, and surface glycoprotein/spike protein of SARS-CoV-2), two human proteins (insulin protein and insulin receptor protein) and food protein (rice Q10MI4). As depicted in Table 6, we does not found any BLAST hits against rice protein, it means that it does not activate/induce TNF-α production. This strategy can be used to scan TNF-α inducing regions in other foods or Genetically modified (GM) foods. Similarly, in the case of human insulin receptor protein, we do not found any hits. Interestingly, we discovered that human insulin hormone which is a small protein contains highest percentage of TNF-α inducing regions i.e., 55.21% (See Table 6). Which completely shows that elevation in insulin levels is responsible for the production of TNF-α peptides/epitopes. This observation agree with the previous studies where they have demonstrated that insulin resistance patients have higher levels of TNF-α [42, 43].

**Table 5:** Performance of hybrid model on mouse main and alternative independent datasets.

| E-value | Main Dataset |  |  |  |  |  | Alternate Dataset |  |  |  |  |  |
| --- | --- | --- | --- | --- | --- | --- | --- | --- | --- | --- | --- | --- |
|  | Sens | Spec | Acc | AUROC | AUPRC | MCC | Sens | Spec | Acc | AUROC | AUPRC | MCC |
| 1.00E-06 | 61.68 | 59.81 | 60.75 | 0.64 | 0.61 | 0.22 | 65.42 | 65.42 | 65.42 | 0.73 | 0.74 | 0.31 |
| 1.00E-05 | 69.16 | 52.34 | 60.75 | 0.63 | 0.61 | 0.22 | 64.49 | 68.22 | 66.36 | 0.73 | 0.75 | 0.33 |
| 1.00E-04 | 58.88 | 59.81 | 59.35 | 0.64 | 0.61 | 0.19 | 66.36 | 66.36 | 66.36 | 0.73 | 0.74 | 0.33 |
| 1.00E-03 | 62.62 | 64.49 | 63.55 | 0.67 | 0.65 | 0.27 | 66.36 | 67.29 | 66.82 | 0.73 | 0.74 | 0.34 |
| 1.00E-02 | 61.68 | 62.62 | 62.15 | 0.68 | 0.66 | 0.24 | 67.29 | 65.42 | 66.36 | 0.75 | 0.79 | 0.33 |
| 1.00E-01 | 62.62 | 65.42 | 64.02 | 0.70 | 0.69 | 0.28 | 66.36 | 67.29 | 66.82 | 0.77 | 0.81 | 0.34 |
| 1.00E+00 | 61.68 | 62.62 | 62.15 | 0.66 | 0.64 | 0.24 | 68.22 | 69.16 | 68.69 | 0.76 | 0.78 | 0.37 |
| 1.00E+01 | 60.75 | 60.75 | 60.75 | 0.66 | 0.64 | 0.22 | 65.42 | 65.42 | 65.42 | 0.71 | 0.74 | 0.31 |
| 1.00E+02 | 60.75 | 60.75 | 60.75 | 0.65 | 0.64 | 0.22 | 66.36 | 65.42 | 65.89 | 0.71 | 0.74 | 0.32 |

**Table 6:**
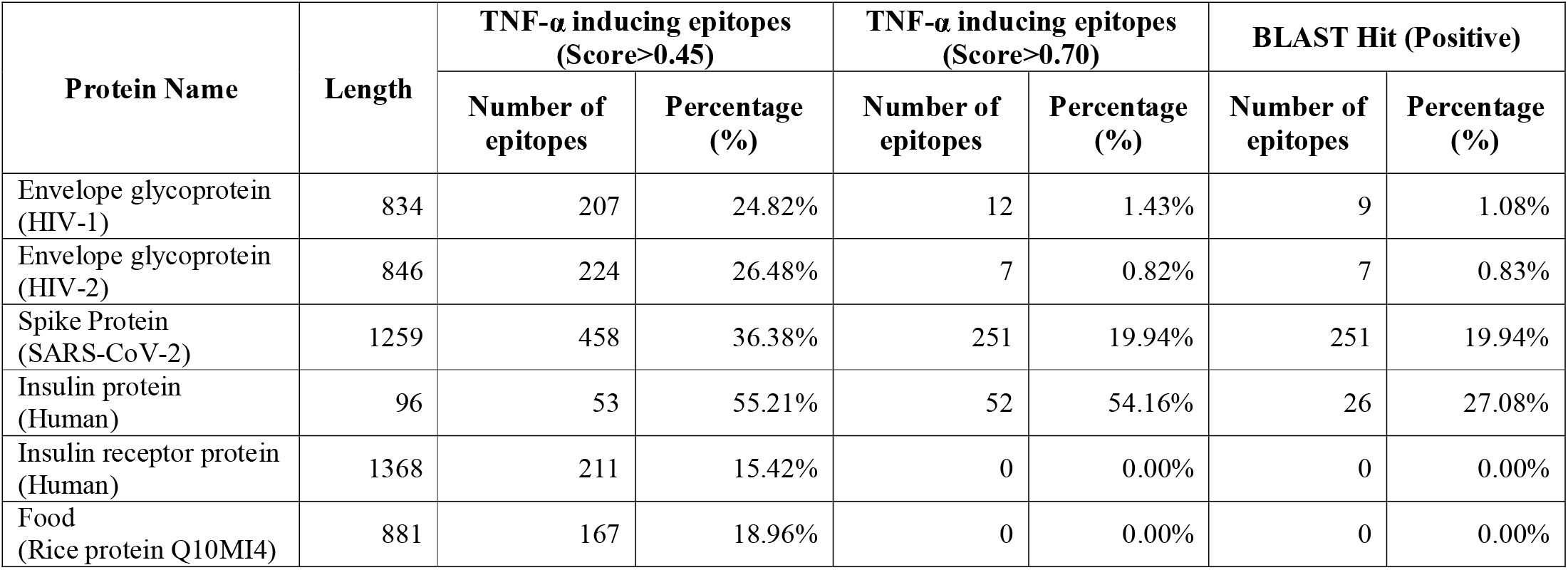
Potential TNF-α inducing epitopes predicted by Protein Scan module of TNFepitope server in 3 viral proteins (HIV-1, HIV-2, and SARS-CoV-2), 2 human proteins (insulin and insulin receptor) and 1 food protein (rice Q10MI4).

In addition, various studies have reported that elevated levels of TNF-α is associated with the pathogenesis of viral infections such as (human immunodeficiency virus (HIV) and SARS-CoV-2) [20, 44–46]. As shown in Table 6, the envelope proteins of HIV-1 and HIV-2 possesses 24.82% and 26.48% TNF-α inducing regions, while the spike protein of SARS-CoV-2 have 36.38% TNF-α inducers, which supports the previous studies where severity in COVID-19 patients is associated with the high levels of TNF-α. In Supplementary Table S7, we have provided the top-most TNF-α inducing epitopes of HIV-1, HIV-2, spike protein and human insulin protein. The complete results for each protein in provided in Supplementary Table S8-S13. These results indicates that our study can be used to measure the levels of TNF-α in different viruses. We hope our findings anticipate the scientific community, working in the era of subunit vaccine designing against deadly viruses and other autoimmune diseases that can be proliferated by the elevation of TNF-α.

## Discussion and Conclusion

Major histocompatibity complex region encodes numbers of proteins including human leukocyte antigen (HLAs) which are necessary for self-recognition, cytokine genes like TNF, LTA, LTB which are responsible for the inflammations [47]. TNF-α is an important inflammatory cytokine released by T cells or macrophages and control a number of signalling pathways within the immune cells; leads to necrosis or cell death [3, 4]. These pathways result in a range of biological responses, such as cell proliferation, differentiation, and survival. TNF-α cytokine employed for cancer treatment and perform anti-cancer activities by inducing inflammation, immune response, and tumor cell apoptosis [48–50]. However, improper and excessive activation of TNF signalling pathway may results into the emergence of pathological diseases such as HIV-I, anorexia, cachexia, obesity, autoimmune disorders including rheumatoid arthritis, diabetes, inflammatory bowel disease, and Crohn’s diseases [51–59]. Several TNF-α inhibitors such as infliximab, etanercept, golimumab, and certolizumab and adalimumab have been developed and approved for clinical use to cure diseases which are associated with abnormal/excessive TNF-α secretion [54, 60].

Mortaz et. al., also report the higher level of soluble TNF-α in the patients of COVID-19 in comparison with the healthy control [61]. Therefore, it is crucial to check for the existence of TNF-α inducing epitopes or to use anti-TNF therapy in a variety of diseases. In the current study, we have attempted to understand the nature of TNF-α inducing peptides and built a prediction model to recognize the epitopes which can induce TNF-α secretion. Dataset play major role in developing machine learning models, hence we have collected experimentally validated TNF-α inducing and non-inducing peptides for human and mouse. In case of alternate negative dataset we have generated random peptides using Swiss-Prot database. To investigate the composition and positional preference, sequence logo and compositional analytical analysis were conducted. We found that TNF-α inducing epitopes are rich in the amino acid residue (L) in human and (N) in mouse datasets. Then after, we employed ‘Pfeature’ to compute 15 types of compositional features using the standalone package.

We have used a number of machine-learning classifiers in order to develop prediction models. Our results indicate that di-peptide composition based features performed best in the case of main and alternate datasets for both human and mouse models. Using DPC based features we have achieved highest AUROC of 0.79 and 0.74 on the human and mouse independent dataset. Of note, our hybrid model (BLAST + machine learning) outperformed others with an AUROC of 0.83 and 0.77 on the human and mouse independent dataset. We have used the best models and created a web server ‘TNFepitope’ for the scientific community, along with a standalone package. TNFepitope (https://webs.iiitd.edu.in/raghava/tnfepitope) is publicly accessible and provide facilities to predict, design, and scan the TNF-α inducing regions. In addition, we have used the ‘Scan’ module of TNFepitope server for the prediction of TNF-α inducing epitopes in the spike protein of SARS-CoV-2, envelope protein (HIV-1 and HIV-2), insulin protein, insulin protein receptor of human and rice protein. We observed higher percentage of TNF-α inducing regions in human insulin protein, followed by spike protein of SARS-CoV-2 and envelope protein (HIV-1 and HIV-2). We believe that this work will be helpful for the researchers in the development of computer-aided vaccine design and enabling them to create subunit vaccines that elicit the appropriate immune response against several TNF-α associated diseases.

## Authors’ contributions

AD and GPSR collected and processed the datasets. AD, SP, KN and GPSR implemented the algorithms and developed the prediction models. AD, SP and GPSR analysed the results. SC, AD and SP created the web server. AD, SJ, SP and SC and GPSR penned the manuscript. GPSR conceived and coordinated the project. All authors have read and approved the final manuscript.

## Acknowledgements

Authors are thankful to the Department of Bio-Technology (DBT) and Department of Science and Technology (DST-INSPIRE) for fellowships and the financial support and Department of Computational Biology, IIITD New Delhi for infrastructure and facilities.

